# Prior expectations in visual speed perception predict encoding characteristics of neurons in area MT

**DOI:** 10.1101/2021.08.23.457358

**Authors:** Ling-Qi Zhang, Alan A. Stocker

**Author notes:** Correspondence: Dr. Alan A. Stocker, Computational Perception and Cognition Laboratory, 421 Goddard Laboratories, 3710 Hamilton Walk, Philadelphia, PA 19104, U.S.A.

## Abstract

Bayesian inference provides an elegant theoretical framework for understanding the characteristic biases and discrimination thresholds in visual speed perception. However, the framework is difficult to validate due to its flexibility and the fact that suitable constraints on the structure of the sensory uncertainty have been missing. Here, we demonstrate that a Bayesian observer model constrained by efficient coding not only well explains human visual speed perception but also provides an accurate quantitative account of the tuning characteristics of neurons known for representing visual speed. Specifically, we found that the population coding accuracy for visual speed in area MT (“neural prior”) is precisely predicted by the power-law, slow-speed prior extracted from fitting the Bayesian model to psychophysical data (“behavioral prior”) to the point that the two priors are indistinguishable in a cross-validation model comparison. Our results demonstrate a quantitative validation of the Bayesian observer model constrained by efficient coding at both the behavioral and neural levels.

**Significance Statement:** Statistical regularities of the environment play an important role in shaping both neural representations and perceptual behavior. Most previous works addressed these two aspects independently. Here we present a quantitative validation of a theoretical framework that makes joint predictions for neural coding and behavior, based on the assumption that neural representations of sensory information are efficient but also optimally used in generating a percept. Specifically, we demonstrate that the neural tuning characteristics for visual speed in brain area MT are precisely predicted by the statistical prior expectations extracted from psychophysical data. As such, our results provide a normative link between perceptual behavior and the neural representation of sensory information in the brain.

## Introduction

Human perception of visual speed is typically biased and depends on stimulus attributes other than the actual motion of the stimulus. Contrast, for example, strongly affects perceived stimulus speed such that a low-contrast drifting grating typically appears to move slower than a high-contrast grating (Thompson, 1982; Stone and Thompson, 1992; Blakemore and Snowden, 1999; Stocker and Simoncelli, 2006). These biases and perceptual distortions are qualitatively consistent with a Bayesian observer that combines noisy sensory measurements with a prior preference for lower speeds (Simoncelli, 1993; Weiss et al., 2002; Stocker, 2006). Previous work has also shown that by embedding the Bayesian observer within a two-alternative forced choice (2AFC) decision process one can “reverse-engineer” the noise characteristics (i.e., likelihood) and prior expectations of individual human subjects from their behavior in a speed-discrimination task (Stocker and Simoncelli, 2004, 2006). This provided both a quantitative validation of the Bayesian observer model and a normative interpretation of human behavior in visual speed perception tasks, which has been confirmed in various later studies (e.g. Welchman et al., 2008; Hedges et al., 2011; Sotiropoulos et al., 2014; Jogan and Stocker, 2015). However, recovering the parameters of a Bayesian observer model from behavioral data is typically difficult due to the intrinsic non-specificity of its probabilistic formulation, which has been grounds for a critical view of the Bayesian modeling approach altogether (Jones and Love, 2011; Bowers and Davis, 2012). The reverse-engineered speed priors in previous studies indeed all showed large variations across subjects, indicating a potential case of over-fitting due to insufficient model constraints (Stocker and Simoncelli, 2006; Hedges et al., 2011; Sotiropoulos et al., 2014; Jogan and Stocker, 2015).

In this article, we show how we addressed this potential problem by developing and validating a tightly constrained Bayesian observer model. We followed a recent proposal to use efficient coding as a constraint that links the likelihood function and the prior expectations of a Bayesian observer (Wei and Stocker, 2012, 2015). The efficient coding hypothesis posits that biological neural systems allocate their limited coding capacity such that overall information transmission is optimized given the stimulus distribution in the natural environment (Barlow, 1961; Laughlin, 1981). It thus establishes a direct relationship between the stimulus distribution and the accuracy of neural representations in sensory systems (Linsker, 1988; McDonnell and Stocks, 2008; Wang et al., 2012; Ganguli and Simoncelli, 2014; Yerxa et al., 2020; Roy et al., 2021). Wei and Stocker (2015) showed how to formulate efficient coding as an information constraint that can be embedded within the probabilistic language of the Bayesian framework. The resulting Bayesian observer model has proven to account for a wide range of phenomena in perception including repulsive biases in perceived visual orientation (Wei and Stocker, 2015; Taylor and Bays, 2018) and the lawful relationship between perceptual bias and discrimination threshold (Wei and Stocker, 2017), but also in more cognitive domains such as subjective preferences judgments (Polania et al., 2019) or the representation of numbers (Cheyette and Piantadosi, 2020; Prat-Carrabin and Woodford, 2021).

The overall goal of our current work was two-fold. First, we aimed for a quantitative validation of this new Bayesian observer model in the domain of visual speed perception. We fit the model to speed discrimination data collected by Stocker and Simoncelli (2006). We found that compared to the model in this original study, the new model allowed us to reverse-engineer much more reliable and consistent estimates of subjects’ prior beliefs while still accurately accounting for subjects’ psychophysical behavior. Second, based on the efficient coding hypothesis we wanted to test whether the reverse-engineered prior expectations are mirrored in the population encoding characteristics of neurons in the motion-sensitive area in the primate brain. The medial temporal (MT) area is widely recognized as the cortical area in the primate brain that selectively represents direction and speed of moving visual stimuli (Zeki, 1974; Newsome and Pare, 1988; Britten et al., 1993; Movshon and Newsome, 1996; Priebe et al., 2003). By analyzing single-cell recordings of a large population of MT neurons (Nover et al., 2005), we found that the sensitivity with which visual speed is encoded in this population (“neural prior”) is precisely predicted by the prior beliefs extracted from the psychophysical data (“behavioral prior”). Our results provide an important quantitative validation of the Bayesian observer model constrained by efficient coding at both the behavioral and neural levels.

## Materials and Methods

### Behavioral prior: Bayesian observer model constrained by efficient coding

#### Data

We reanalyzed the two-alternative-forced-choice (2AFC) speed discrimination data from Stocker and Simoncelli (2006). In each trial of the experiment a subject was shown a pair of horizontally drifting gratings (reference and test), and was asked to choose which one of them was moving faster. The reference grating had one of two contrast levels [0.075, 0.5] and one of six different drifting speeds [0.5, 1, 2, 4, 8, 12] deg/s. The test grating had one of seven different contrast levels [0.05, 0.075, 0.1, 0.2, 0.4, 0.5, 0.8] and its speed was determined by an adaptive staircase procedure (one-up/one-down). There were 72 different individual conditions (i.e., psychometric curves) and each condition contained 80 trials, resulting in a total of 5,760 trials. We excluded one subject (labeled as Subject 3 in Stocker and Simoncelli (2006)) that was only tested at two contrasts and two test speed levels. While we were able to recover a prior from this subject that was highly consistent with the rest of the subjects, it was not possible to perform a meaningful model comparison and cross-validation due to the low number of trials.

#### Model formulation

We use the Bayesian observer model by Wei and Stocker (2015) and embed it within a decision process to predict the binary judgments in the 2AFC experiment (Stocker and Simoncelli, 2006).

Specifically, we assume that “encoding” of the stimulus is governed by an efficient coding constraint such that encoding accuracy, measured as the square-root of Fisher Information (FI), is proportional to the stimulus prior (Brunel and Nadal, 1998; McDonnell and Stocks, 2008; Wei and Stocker, 2016), that is

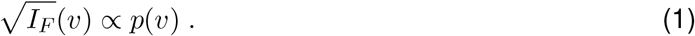

Encoding is described as the conditional probability distribution *p*(*m*|*v*). It determines how stimulus speed *v* is transformed probabilistically into a noisy sensory measurement *m*. We can satisfy the efficient coding constraint (Eq. (1)) by assuming the encoding distribution

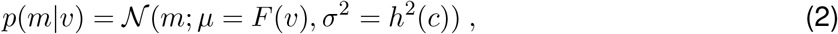

where 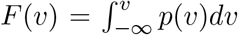 is the cumulative density function (CDF) of *v*. We parameterized the speed prior distribution as the modified power-law function

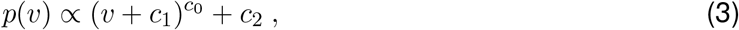

where *c*_0,1,2_ are free and unconstrained parameters. We also tested alternative parameterizations (Fig. 4). The scalar *h*(*c*) determines the amount of total encoding resources (i.e., the overall magnitude of internal noise) at different contrast levels. It can be shown that

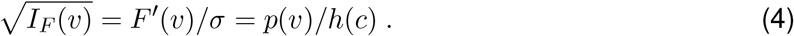

The total amount of encoding resource is measured by 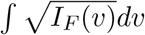 which evaluates to 1*/h*(*c*). In order to numerically handle the unbounded nature of a magnitude variable such as speed (compared to a circular variable such as orientation), we added a small constant (2.5 ∗ 10^*−*3^) to *p*(*v*) such that its CDF did not saturate (i.e., *F* (*v*) is not upper bounded by 1).

To decode (i.e., estimate) the stimulus *v* given a particular sensory representation *m*, we first determine the likelihood function

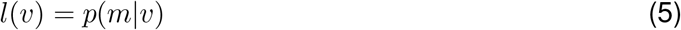

by considering the encoding distribution (Eq. (2)) as a function of *v*. Applying Bayes’ rule and multiplying the likelihood function with the prior *p*(*v*) (Eq. (3)), we then can compute the posterior as

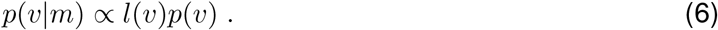

Assuming an *L*_0_ loss function (Stocker and Simoncelli, 2006), the estimate 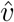 of the stimulus *v* is given as

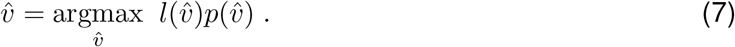

The estimate represents the optimally decoded stimulus 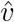 given *m*. It is a deterministic function of *m* (implicit in the likelihood function *l*(*v*)), which we can explicitly express as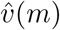. However, *m* is not directly observable in a psychophysical experiment. Thus, we marginalize over *m* to obtain the estimate distribution for a given stimulus *v*,

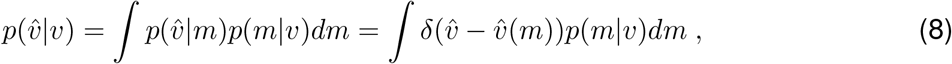

where *δ*(·) is the Dirac delta function.

In a 2AFC speed discrimination experiment, subjects report a binary decision and not a continuous estimate 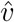. We assume subjects make their choice (i.e., which one is faster) by comparing their estimate 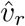 of the reference stimulus with their estimate 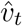 of the test stimulus. Across many repeated trials, these choices follow a binomial distribution with the probability of the test stimulus being perceived faster given as

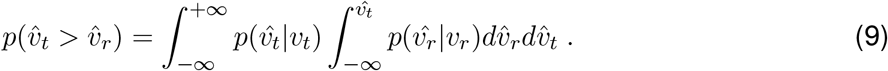

#### Model fitting

If we represent the data in our experiment as *N* triplets (*v*_*ir*_, *v*_*it*_, *k*_*i*_), where *k*_*i*_ ∈ {0, 1} represents the binary choice, then the overall log-likelihood of the model given the data is

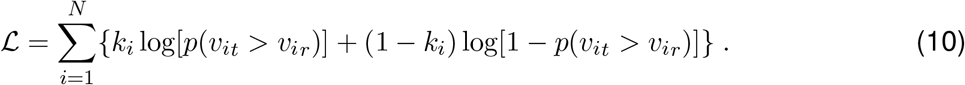

We find the model parameters *c*_0_, *c*_1_, *c*_2_ and *h*(*c*) by maximizing L using MATLAB’s *fminsearchbnd* algorithm. Note that the model is highly constrained: For each subject, we jointly fit a single three-parameter prior distribution plus one scalar noise parameter *h*(*c*) for each of the 7 contrast levels to the data from all 72 conditions.

#### Alternative prior parameterization

In order to assess the consistency and stability of our reverse-engineered prior distributions, we also tested two alternative parameterizations:

- a Gamma distribution: *p*(*v*; *α, β*) ∝ |*v*|^*α−*1^*e*^*−β*|*v*|^
- a piece-wise log-linear function with 18 sample points 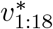 equally distributed in logarithmic space in the range *v* = [0…50] deg/s. Each corresponding 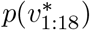 value is a free prior parameter; prior density values are linearly interpolated between those values.

For comparison we also fit a Gaussian prior with *p*(*v*; *σ*^2^) = 𝒩 (*v*; *µ* = 0, *σ*^2^).

#### Weber’s law and power-law prior

With our model, it is possible to analytically predict discrimination threshold Δ_*v*_ and Weber fraction 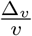 for any given prior distribution. It has been shown that discrimination threshold is inversely proportional to the square-root of FI (Seriès et al., 2009; Wei and Stocker, 2017), thus

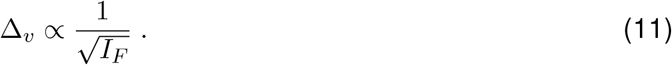

According to the efficient coding constraint (Eq. (1)), we can substitute 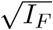 with *p*(*v*) and find

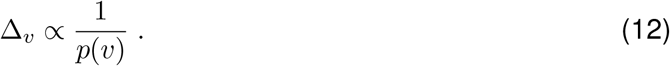

This equation allows us to predict discrimination threshold for any prior density (up to a factor). For the modified power-law prior with exponent *c*_0_ = −1 (Eq. (3)) we find

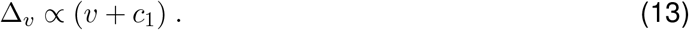

By further setting *c*_1_ = 0, we obtain *v* ∝ Δ_*v*_, which is the definition of Weber’s law (i.e., a constant Weber fraction). For non-zero *c*_1_ the Weber fraction changes to

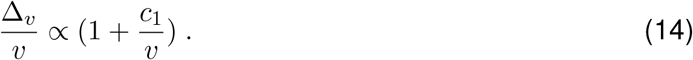

At high speeds, 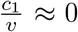 and thus the Weber fraction is constant. At low speeds, 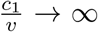 causing 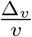 to increase.

Our efficient coding constraint implies that the stimulus is transformed according to the CDF of *v* (Eq. (2)). For a power-law prior with exponent *c*_0_ = −1, the CDF is

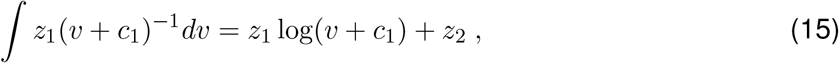

which is precisely the logarithmic transformation that has been previously used for describing the speed tuning of MT neurons (Nover et al., 2005).

### Neural prior: MT encoding analysis

#### Data

We reanalyzed the electrophysiological recording data from Nover et al. (2005). Neurons in area MT of several macaque monkeys were individually identified. Each identified neuron was then tested with a random-dot motion stimulus moving with one of eight speeds [0, 0.5, 1, 2, 4, 8, 16, 32] deg/s. Stimulus location, direction, size and disparity were individually optimized for each neuron. Every stimulus speed was presented three to seven times. We considered the mean firing rate over the entire stimulus duration (1.5 s) as a neuron’s single-trial response. We analyzed a total of 480 neurons.

#### Population Fisher information

Following Nover et al. (2005), we fit each neuron’s mean firing rate as a function of stimulus speed with a Gaussian tuning curve in log-speed

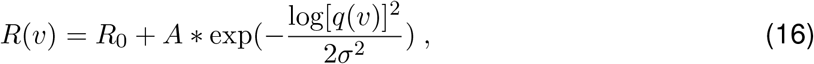

Where 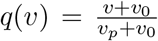. Parameters *R*_0_, *A, σ*^2^, *v*_0_, and *v*_*p*_ are determined by minimizing the sum of squared difference of the observed and predicted firing rates. A maximum-likelihood fit assuming firing rate variability to follow a Poisson distribution produced very similar results.

We computed the population FI for different assumptions about neurons’ response variabilities and their pair-wise noise correlations within the population. First, we assumed that response noise is independent between neurons in the population and response variability is well-described by a Poisson process. In this case, the population FI is calculated as

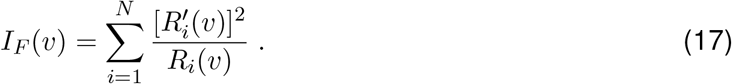

The “neural prior” (the prior that corresponds to the measured MT encoding precision assuming efficient encoding) is then equivalent to the normalized square-root of FI, thus

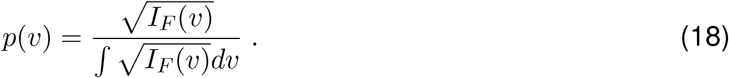

As in Nover et al. (2005), we also repeated the above analysis using an alternative tuning-curve model (Gamma distribution function) and obtained very similar results.

Next, we estimated the population FI by adjusting the Poisson model with an explicit estimate of the Fano factor *F*_*i*_ for each neuron

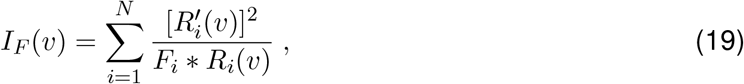

where *F*_*i*_ was obtained by linearly regressing the neuron’s firing rate variance against its firing rate mean. Finally, we computed the linear Fisher information (Kanitscheider et al., 2015; Kohn et al., 2016)

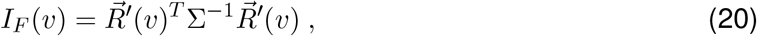

where Σ is the noise correlation matrix and is the identity matrix for the independent noise case. To understand the effect of tuning preference-dependent noise correlations observed in area MT (Huang and Lisberger, 2009), we adopted the limited-range correlation model

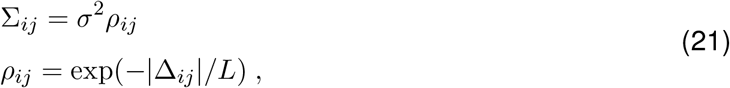

where Δ_*ij*_ is difference in log-speed preference between neuron *i* and neuron *j, σ*^2^ is the noise variance, and *L* is the overall correlation strength (Abbott and Dayan, 1999). For the simulations shown in Fig. 7E, we set *σ*^2^ = 1 as the prior is only determined by the shape of the population FI and *L* = [0.5, 1.0, 2.5] for low, medium and high correlation strengths.

#### Cross-validation

We performed a five-fold cross-validation procedure. The trial data for each condition were first randomly and equally divided into 5 groups. For each group, the model was fit to the data of the remaining four groups (training), and then evaluated on the group’s data (validation). Model validation performance was measured as the log-likelihood of the fit model given the validation data. The entire procedure was repeated 20 times, resulting in 100 estimates of the model validation likelihood. For the behavioral prior condition, we considered the full observer model using the fit prior for that run. For the neural prior condition, we assumed the prior to be fixed and equal to the prior extracted from the population FI analysis with only the contrast-dependent noise parameters being fit on each run. The same procedure was used to compute the validation likelihoods of the original, less constrained Bayesian observer model (Stocker and Simoncelli, 2006), and of individual Weibull fits to every condition. Log-likelihood values in Fig. 8B were normalized to the range set by a lower bound given by the log-likelihoods of a coin-flip model for the decision (i.e. a model with a fixed decision probability of 0.5), and an upper bound determined by the values of the Weibull fits.

### Code Accessibility

Data and analysis code are available through GitHub: https://github.com/cpc-lab-stocker/Speed_Prior_2021

## Results

We model speed perception as an efficient encoding, Bayesian decoding process (Fig. 1A). On any given trial the speed *v* of a visual stimulus is represented by a noisy and bandwidth-limited sensory measurement *m*. Following Wei and Stocker (2012, 2015), we assume that “encoding” of the stimulus is governed by an efficient coding constraint (Eq. (1)) such that encoding accuracy, measured as the square-root of Fisher Information (FI), is proportional to the stimulus prior *p*(*v*) (Brunel and Nadal, 1998; McDonnell and Stocks, 2008; Wei and Stocker, 2016). This constraint promotes a more accurate encoding of speeds for which the prior density is high. It determines the observer’s uncertainty about the actual stimulus speed given a particular sensory measurement (i.e., the likelihood function *p*(*m*|*v*)). For “decoding”, this likelihood function *p*(*m*|*v*) is combined with the stimulus prior *p*(*v*), resulting in the posterior *p*(*v*|*m*). Lastly, a percept 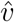 (i.e., an estimate) is computed based on the posterior and a loss function.

**Figure 1:**
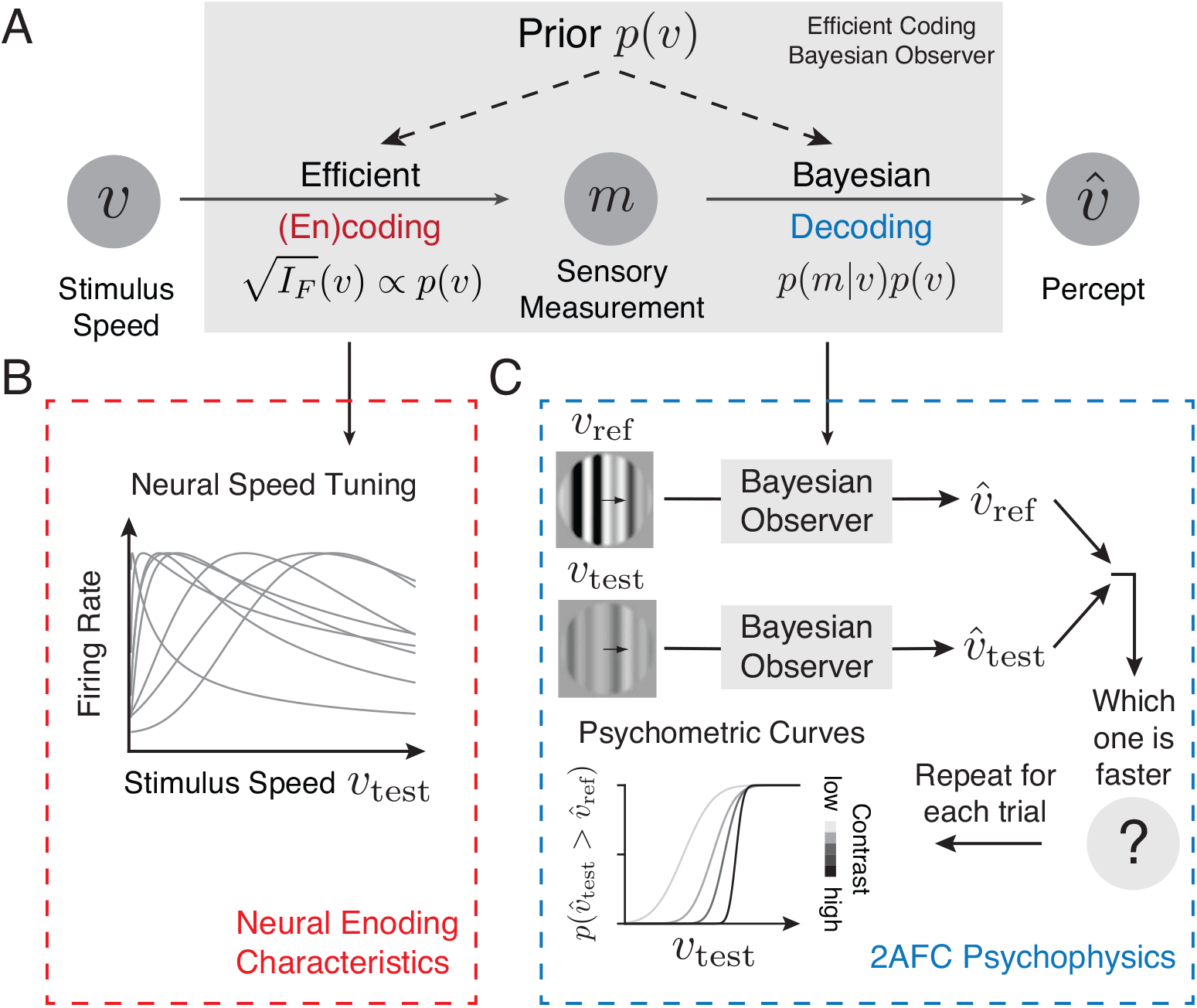
Bayesian observer model constrained by efficient coding. **A)** We model speed perception as an efficient encoding, Bayesian decoding process (Wei and Stocker, 2012, 2015). Stimulus speed *v* is encoded in a noisy and resource-limited sensory measurement *m* with an encoding accuracy that is determined by the stimulus prior *p*(*v*) via the efficient coding constraint (Eq. (1)). Ultimately, a percept is formed through a Bayesian decoding process that combines the likelihood *p*(*m*|*v*) and prior *p*(*v*) to compute the posterior *p*(*v*|*m*), and then selects the optimal estimate 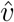 according to a loss function. Encoding and decoding are linked and jointly determined by the prior distribution over speed. **B)** Efficient coding determines the accuracy of the neural representation of visual speed (i.e., the tuning characteristics of neurons in area MT). **C)** Embedding the Bayesian observer within a decision process provides a model to predict psychophysical behavior in a two-alternative forced choice (2AFC) speed discrimination task.

A unique feature of the new model is that the stimulus distribution jointly determines encoding and decoding of the Bayesian observer model (Wei and Stocker, 2012). Thus, both the encoding characteristics of neurons representing visual speed (Fig. 1B), and the psychophysical behavior of subjects in speed perception (Fig. 1C) should be consistent with the prior belief of the observer about the statistical regularities of visual speed.

### Extracting the “behavioral prior”

We fit our model to the psychophysical speed discrimination data collected by Stocker and Simon-celli (2006). On each trial of their experiment, subjects were shown a pair of horizontally drifting gratings (reference and test stimulus), and were asked to choose which one was moving faster (Fig. 1C). For each combination of reference and test, a full psychometric curve was measured by repeating the trials at different test speeds chosen by an adaptive staircase procedure. A total combination of 72 conditions representing reference and test stimuli at different speeds and contrast levels were tested, resulting in 72 different psychometric functions (see *Methods* and Stocker and Simoncelli (2006) for details).

In contrast to the original model (Stocker and Simoncelli, 2006), the new observer model directly links the likelihood function and the prior distribution (Wei and Stocker, 2012). Thus perceived speed is determined by subjects’ prior expectations and a contrast-dependent internal noise parameter that reflects the total amount of represented sensory information (Wei and Stocker, 2015, 2016). Our goal was to find the prior distribution *p*(*v*) and the noise parameters *h*(*c*) that best accounted for subjects’ individual perceptual behavior. In order to fit the observer model, we embedded it within a binary decision process (Fig. 1C). On each trial, speed estimates for both the reference and the test stimuli are performed, and then subjects are assumed to respond according to which estimate is faster. Entire psychometric functions are predicted by marginalizing over the (unobserved) sensory measurement (*Methods*).

We jointly fit our model for every subject to all 72 conditions using a maximum-likelihood procedure. The free parameters of the model consisted of a parametric description of the prior and one noise parameter for each stimulus contrast. Following previous studies (Stocker and Simoncelli, 2006; Hedges et al., 2011; Jogan and Stocker, 2015), we parameterized the prior distribution as a modified power-law function (Eq. (3)). Figure 2A shows the data and model fit for a few example conditions for exemplary Subject 1. Fit parameter values for all subjects are listed in Table 1. Overall, the model predicts psychometric curves that are similar to those obtained from fitting a Weibull function. The log-likelihood of the new model is close to that of separate Weibull fits to every individual condition (Fig. 2B). Figure 2C further illustrates that the new model performs as well as the original, less constrained Bayesian observer model (Stocker and Simoncelli, 2006). A more detailed model comparison utilizing cross-validation is provided in a later section.

**Table 1:** Fit parameter values of the prior density 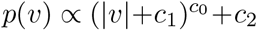 and the contrast-dependent noise *s* = *h*(*c*) for every subject.

| Subject | $c_0$ | $c_1$ | $c_2$ | $h(0.05)$ | $h(0.075)$ | $h(0.10)$ | $h(0.20)$ | $h(0.40)$ | $h(0.50)$ | $h(0.80)$ |
| --- | --- | --- | --- | --- | --- | --- | --- | --- | --- | --- |
| 1 | -0.790 | 0.003 | $6 * 10^{-5}$ | 0.035 | 0.027 | 0.028 | 0.019 | 0.016 | 0.013 | 0.012 |
| 2 | -0.867 | 0.002 | $10^{-8}$ | 0.045 | 0.041 | 0.038 | 0.030 | 0.023 | 0.020 | 0.010 |
| 3 | -1.045 | 0.220 | $1 * 10^{-5}$ | 0.061 | 0.045 | 0.041 | 0.029 | 0.017 | 0.019 | 0.026 |
| 4 | -1.097 | 0.248 | $7 * 10^{-6}$ | 0.043 | 0.040 | 0.036 | 0.028 | 0.018 | 0.017 | 0.016 |

**Figure 2:**
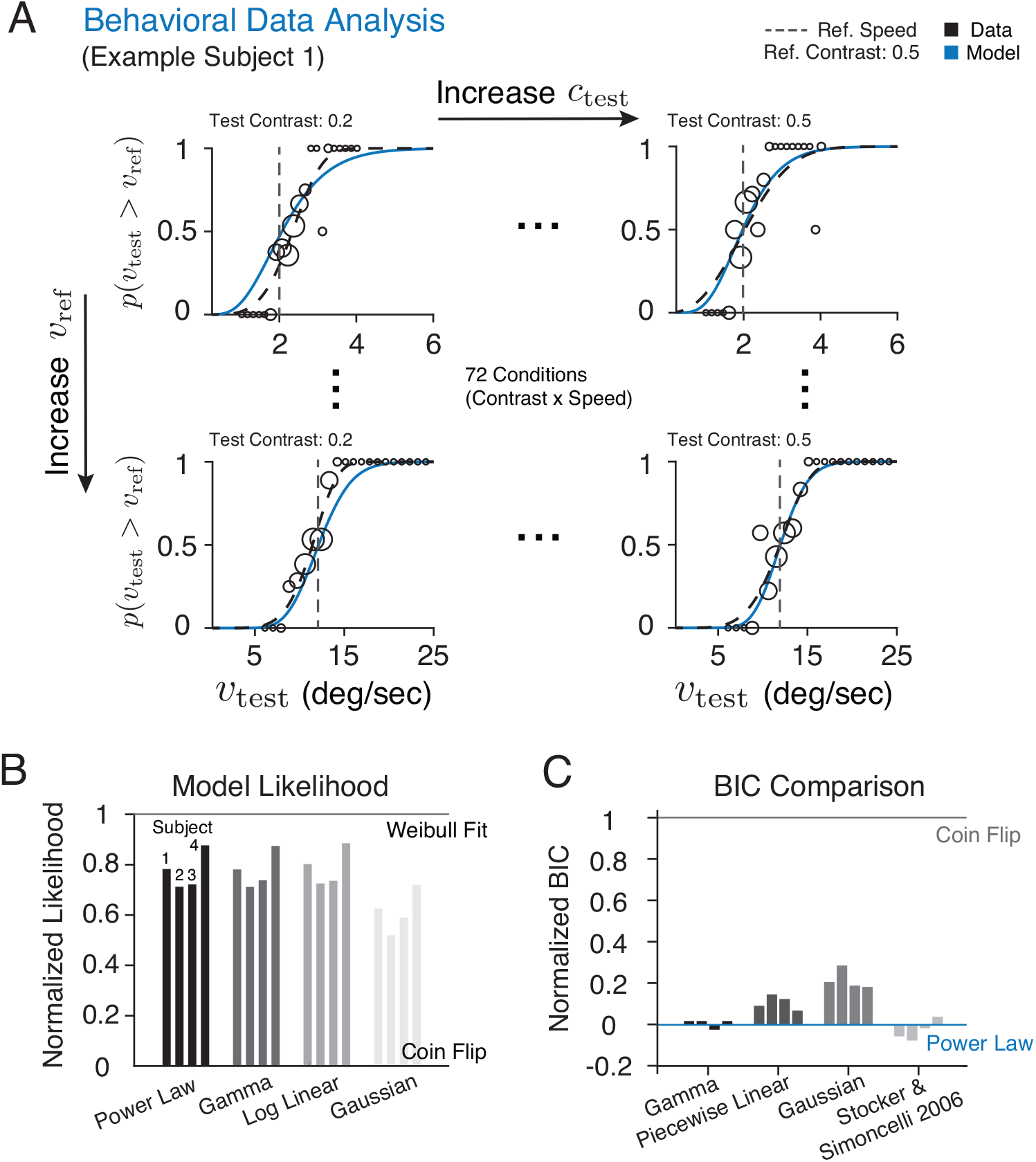
Extracting the “behavioral prior”. **A)** We jointly fit the Bayesian observer model to psychophysical data across all contrast and reference speed conditions (72 conditions total). Shown are a few conditions for exemplary Subject 1. Circle sizes are proportional to the number of trials at that test speed. The dashed curves are Weibull fits to each conditions, and the solid blue curves represent the model prediction. See Extended Data Figure 2 - 1 and 2 - 2 for psychometric curves and model fits of all 72 conditions. **B)** Log-likelihood values of the best-fitting model for each subject using four different prior parameterizations including a power-law function, Gamma distribution, piece-wise log-linear function, and a Gaussian distribution, respectively. Values are normalized to the range set by a coin flip model (lower bound) and Weibull fits to individual psychometric curves (upper bound). **C)** The relative Bayesian Information Criterion (BIC) values for the different parameterizations as well as the original, less constrained Bayesian observer model by Stocker and Simoncelli (2006). Values are normalized to the range set by the efficient coding Bayesian model with power-law parameterization and the coin flip model (lower is better). See *Methods* for details.

Importantly, the reverse-engineered prior expectations are much more consistent across subjects than those obtained from using the original model (Stocker and Simoncelli, 2006; Hedges et al., 2011; Jogan and Stocker, 2015; Sotiropoulos et al., 2014). The exponent *c*_0_, for example, is now close to a value of −1 for every subject rather than varying over an order of magnitude (Table 1). Furthermore, values of the contrast-dependent noise parameter monotonically decrease as a function of contrast as expected and are consistent with the functional description of the contrast response curve of cortical neurons (Fig. 3).

**Figure 3:**
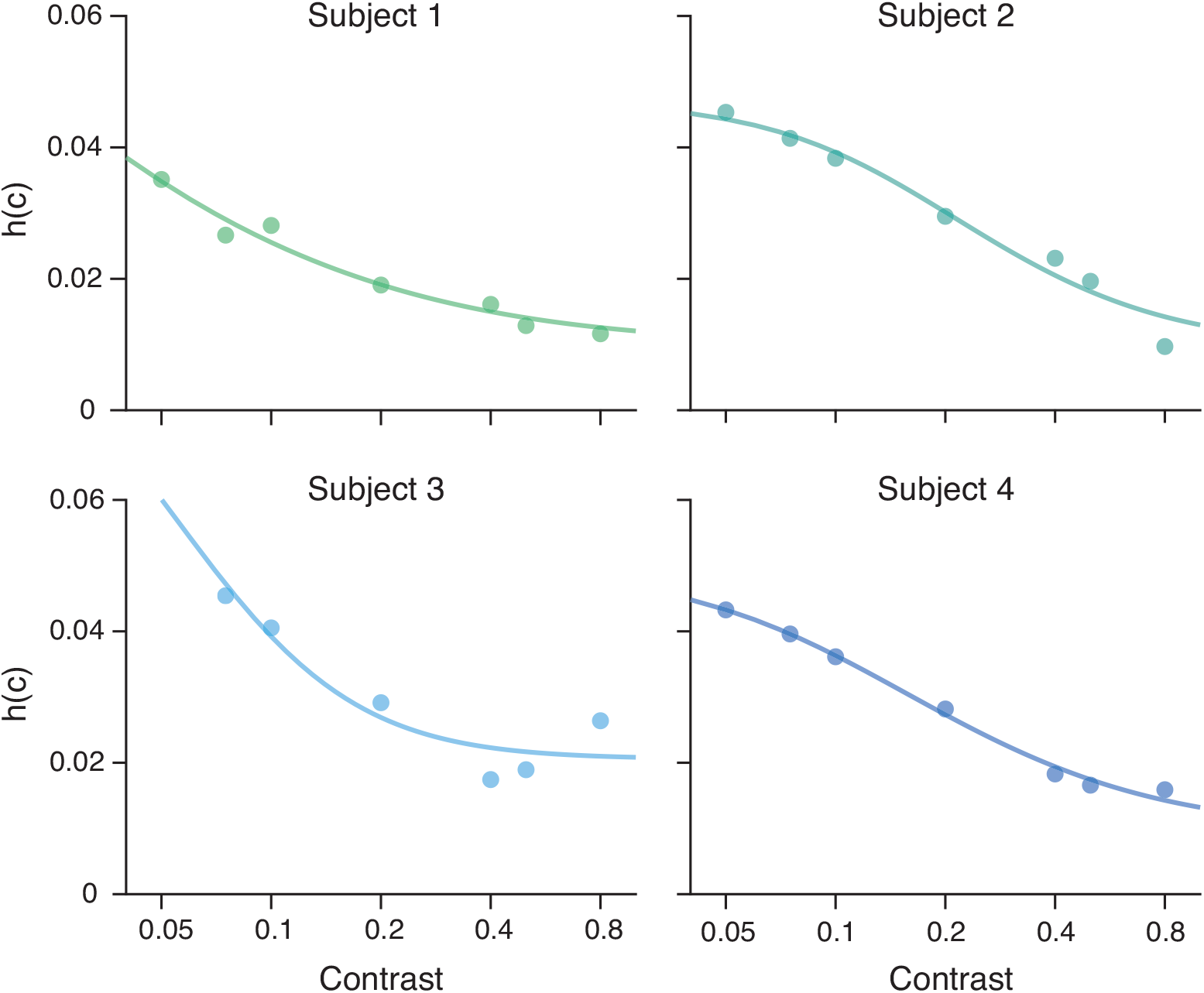
Contrast-dependent noise. Fit parameters values as a function of contrast plotted for every subject. Bold lines represent fits with a parametric description of the contrast response function of cortical neurons 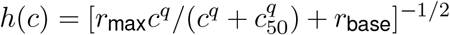 (Sclar et al., 1990; Albrecht and Hamilton, 1982; Heuer and Britten, 2002).

In order to test the impact of choosing a power-law parameterization for the prior distribution (Eq. 3), we performed model fits using two other parameterizations with increasing degrees of freedom (i.e., a Gamma distribution and a piece-wise log-linear function), and also a Gaussian prior for comparison (*Methods*). The model fits well for all but the Gaussian prior, resulting in similar log-likelihood values (Fig. 2B) although the BIC value is higher for the log-linear parameterization due to its large number of parameters (Fig. 2C). Crucially, however, the shapes of the fit prior distributions are very similar across the different parameterizations, all exhibiting a power-law like, slow-speed preferred distribution (Fig. 4). The obvious exception is the Gaussian parameterization because it is unsuited to approximate a power-law function.

**Figure 4:**
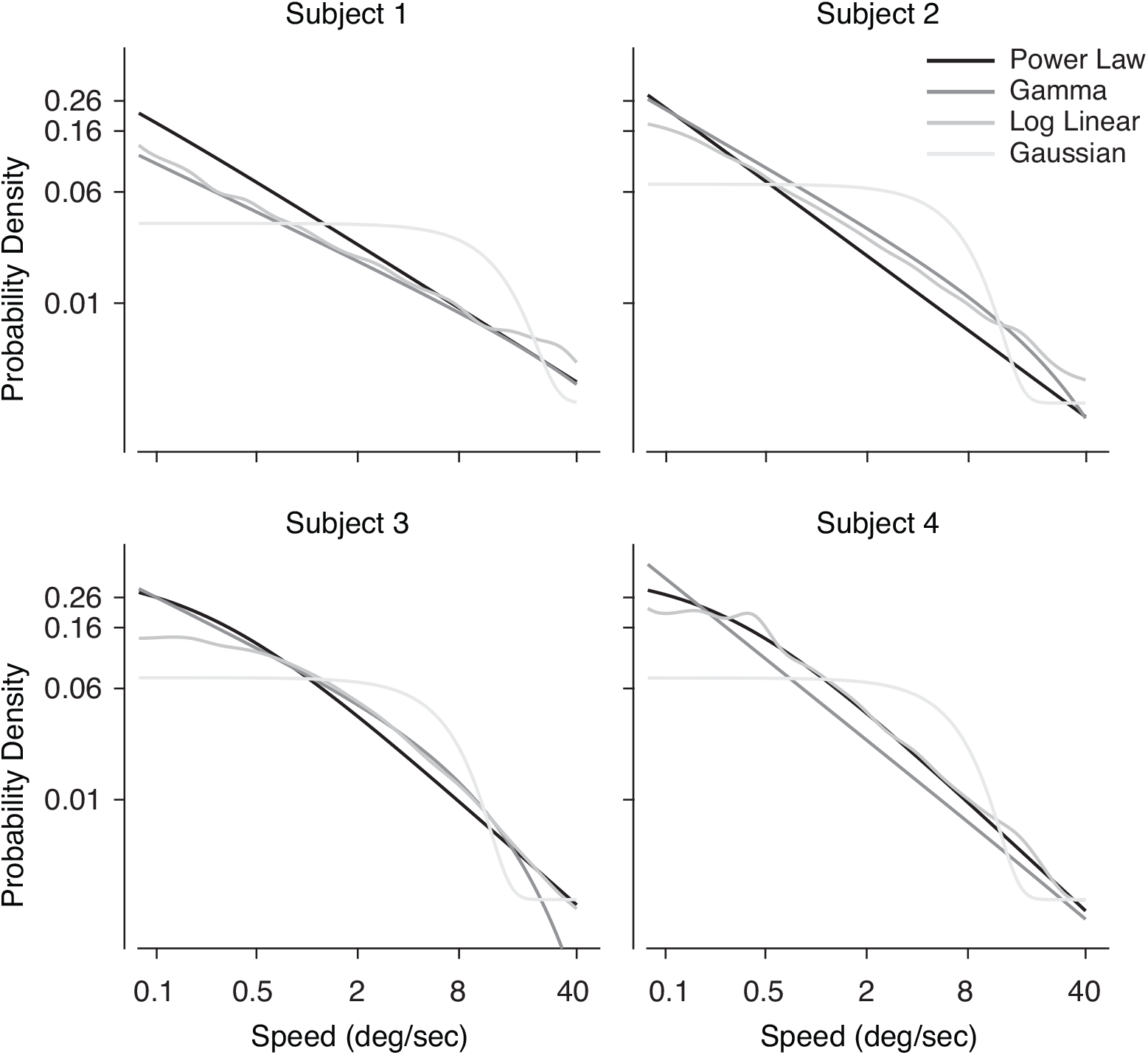
Prior parameterization. Fit prior density functions for each subject using four different parameterizations including a power-law function, a Gamma distribution, a piece-wise log-linear function, and a Gaussian distribution, respectively. Only the Gaussian provides a relative poor fit of the data (see also Fig. 2B).

We further validated our model by comparing its predictions for contrast-induced biases and discrimination thresholds to subjects’ behavior. To quantify bias, we computed the ratio of test speed to reference speed at the point of subjective equality (PSE, defined as the 50% point of the psychometric curve). If a lower-contrast test stimulus is indeed perceived to be slower, then its physical speed will need to be higher in order to match the perceived speed of the higher-contrast reference. Thus, a contrast-induced slow-speed bias is manifested by a PSE ratio greater than 1 when the test contrast is lower than the reference, and vice versa. As shown in Fig. 5A, subjects clearly underestimated the speeds of low-contrast stimuli, an effect that occurred at any contrast level and speed. Furthermore, subjects’ thresholds increase monotonically with speed (Fig. 5B). While they follow Weber’s law at higher speeds, they deviate from a constant Weber-fraction at slow speeds, which is well documented (Stocker and Simoncelli, 2006; McKee et al., 1986; De Bruyn and Orban, 1988). Our new model is able to capture both the contrast-induced slow-speed bias and the discrimination threshold behavior with an accuracy comparable to the original model (Stocker and Simoncelli, 2006) (Fig. 2C).

**Figure 5:**
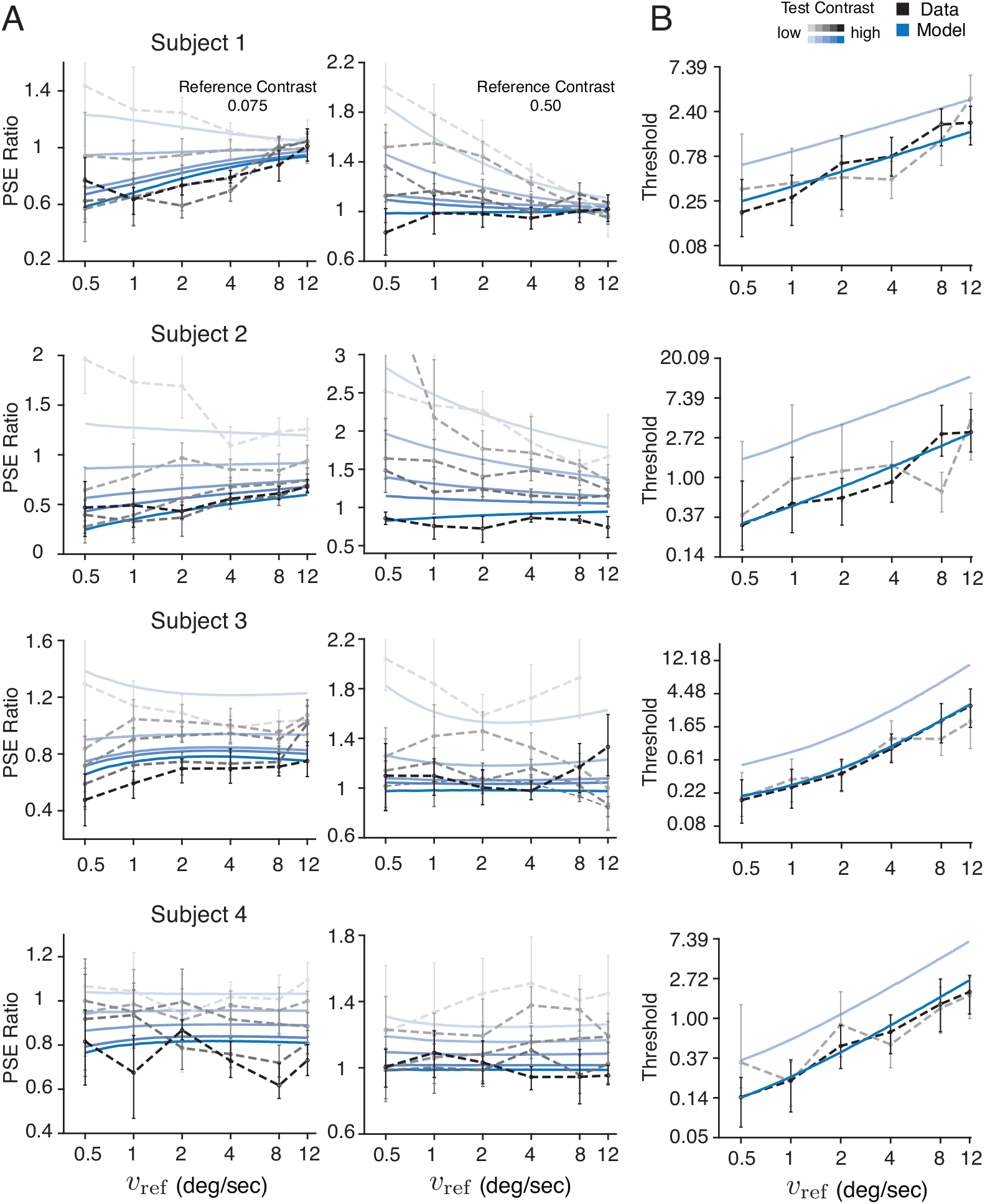
Predicted contrast-dependent bias and discrimination threshold. **A)** Ratios of the test relative to reference speed at the point of subjective equality (PSE) extracted from the Weibull fits (black) and our model (blue). Shading levels correspond to different contrast levels (0.05, 0.1, 0.2, 0.4, 0.8) of the test stimulus (darker means higher contrast). The reference stimulus has a contrast of 0.075 in left column, and 0.5 in the right column. **B)** Speed discrimination thresholds, defined as the difference in stimulus speed at the 50% and 75% points of the psychometric curve, at two different contrast levels (0.075, 0.5), extracted from the Weibull fits (black) and our model (blue). Error bars indicate the 95% confidence interval across 500 bootstrap runs.

Note that the predicted higher threshold values for the low-contrast stimulus condition are not particularly evident in the data (Fig. 5B). Previous studies, however, have convincingly demonstrated that lower stimulus contrasts lead to higher speed discrimination thresholds under various stimulus configurations (Panish, 1988; Turano and Pantle, 1989; Horswill and Plooy, 2008; Champion and Warren, 2017). Thus we believe that the experimental design in Stocker and Simoncelli (2006), in particular the deliberate compromise in choosing a low number of trials per condition (only 80 trials per psychometric curve) in order to test subjects over a large range of different contrast/speed combinations, is responsible for the noisy, overlapping threshold estimates. The ability to obtain reliable threshold estimates was further limited by the use of a stair-case procedure optimized for inferring the PSE rather than the slope of the psychometric curves (see Stocker and Simoncelli (2006) for details). Future investigations will be required to fully resolve this discrepancy.

Despite its constrained nature, the model can well account for individual differences across subjects. Differences in the values of the contrast-dependent noise parameter determine individual variations in bias and threshold magnitude. In addition, when the prior exponent is close to −1, the PSE ratios are mostly constant across different speeds (Fig. 5A; e.g., Subject 3 and 4) (Wei and Stocker, 2017), whereas an exponent larger than −1 predicts relative biases that decrease for higher stimulus speeds (Fig. 5A; e.g., Subject 1).

Lastly, previous models have mainly focused on the contrast-induced speed bias and how it can be attributed to a slow-speed prior that shifts the percept toward slower speeds for increasing levels of sensory uncertainty (Weiss et al., 2002; Hürlimann et al., 2002; Stocker, 2006; Stocker and Simoncelli, 2006; Hedges et al., 2011; Rokers et al., 2018; Lakshminarasimhan et al., 2018). While this is still the case for our new model, the monotonic increase in threshold is now also a direct consequence of the slow-speed prior: Since higher speeds are less likely, efficient coding dictates that less neural resources are allocated for their representation resulting in a larger threshold. In fact, the predicted Weber fractions based on subjects’ reverse-engineered priors closely resemble previous psychophysical measurements (Fig. 6; *Methods*).

**Figure 6:**
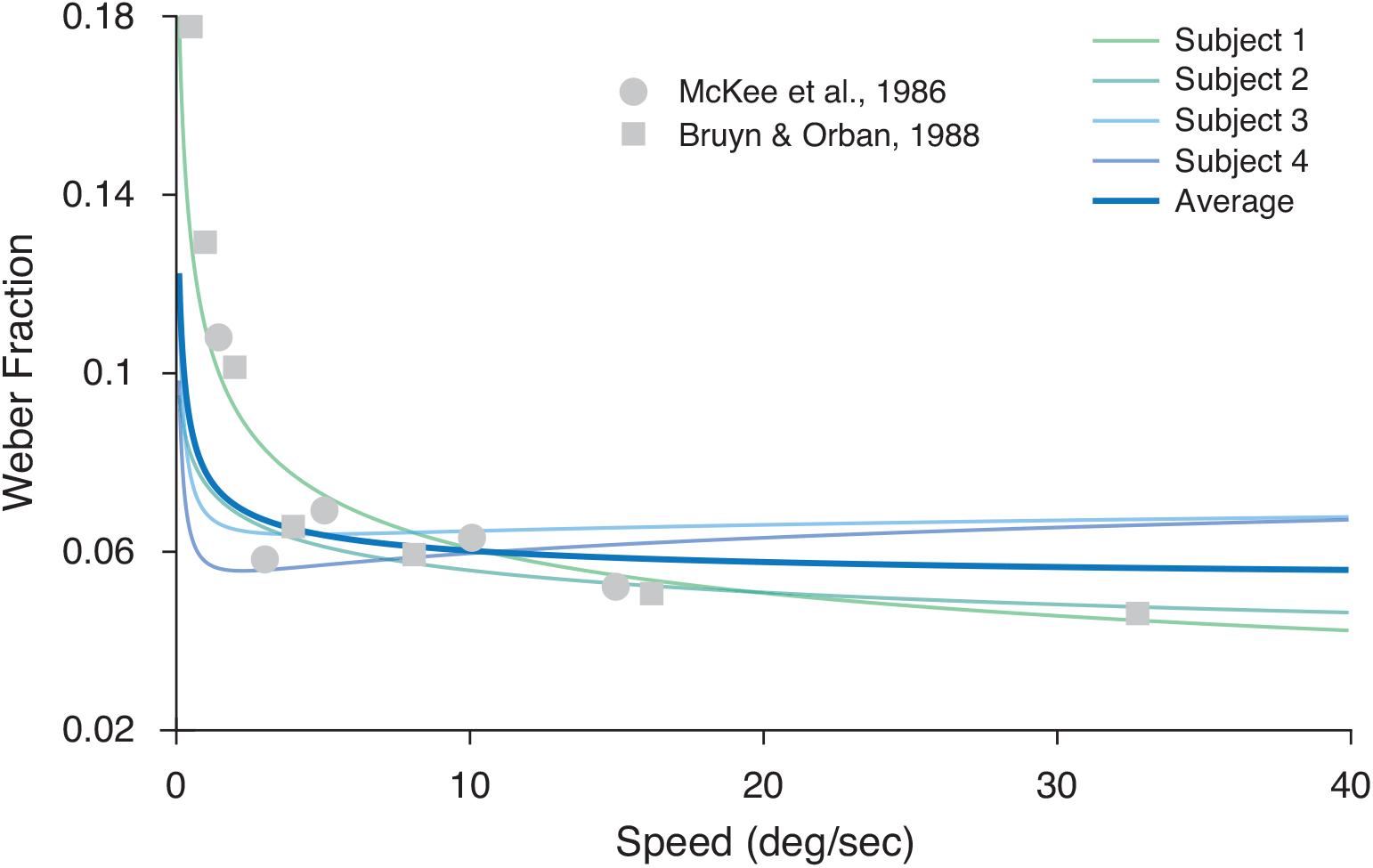
Predicted Weber fraction. Predicted Weber fractions Δ*v/v* based on the reverse-engineered behavioral priors of the four individual and the average subject are shown in comparison to previously reported psychophysical measurements (McKee et al., 1986; De Bruyn and Orban, 1988). In fact, we can analytically show that the modified power-law prior with an exponent of *c*_0_ = −1 will predict both the constant Weber fraction at higher speed and its deviation at slow speeds (*Methods*). Note that since the Weber fraction is predicted up to a factor, the predictions are scaled to the level of the data. De Bruyn and Orban (1988) also found deviations from Weber’s law at extremely high speeds (256 cycles/deg) which is not depicted here.

### Extracting the “neural prior”

The efficient coding constraint of the model predicts that the neural encoding of visual speed reflects the stimulus prior distribution (Wei and Stocker, 2012, 2016; Ganguli and Simoncelli, 2014). Thus, if our new model is correct, then the reverse-engineered “behavioral prior” should be a good predictor of the neural encoding characteristics of visual speed. Specifically, we expect the neural encoding accuracy, measured as the square-root of the neural population FI, to match the extracted speed prior (Eq. (1)).

To test this prediction, we analyzed the encoding characteristics of a large population of neurons in area MT. Our analysis was based on single-cell recorded data from the macaque brain (Nover et al., 2005). The data contained repeated spike counts from 480 MT neurons responding to random dot motion stimuli moving at eight different speeds. Following the original study (Nover et al., 2005), we fit a log-normal speed tuning curve model for each neuron in the data set (Fig. 7A). This tuning curve model accurately described the mean firing rates of the majority of the neurons (Fig. 7B). Under the assumption that neural response variability was well captured by a Poisson distribution, it is then straight-forward to compute the FI of individual neurons (Fig. 7A; *Methods*).

**Figure 7:**
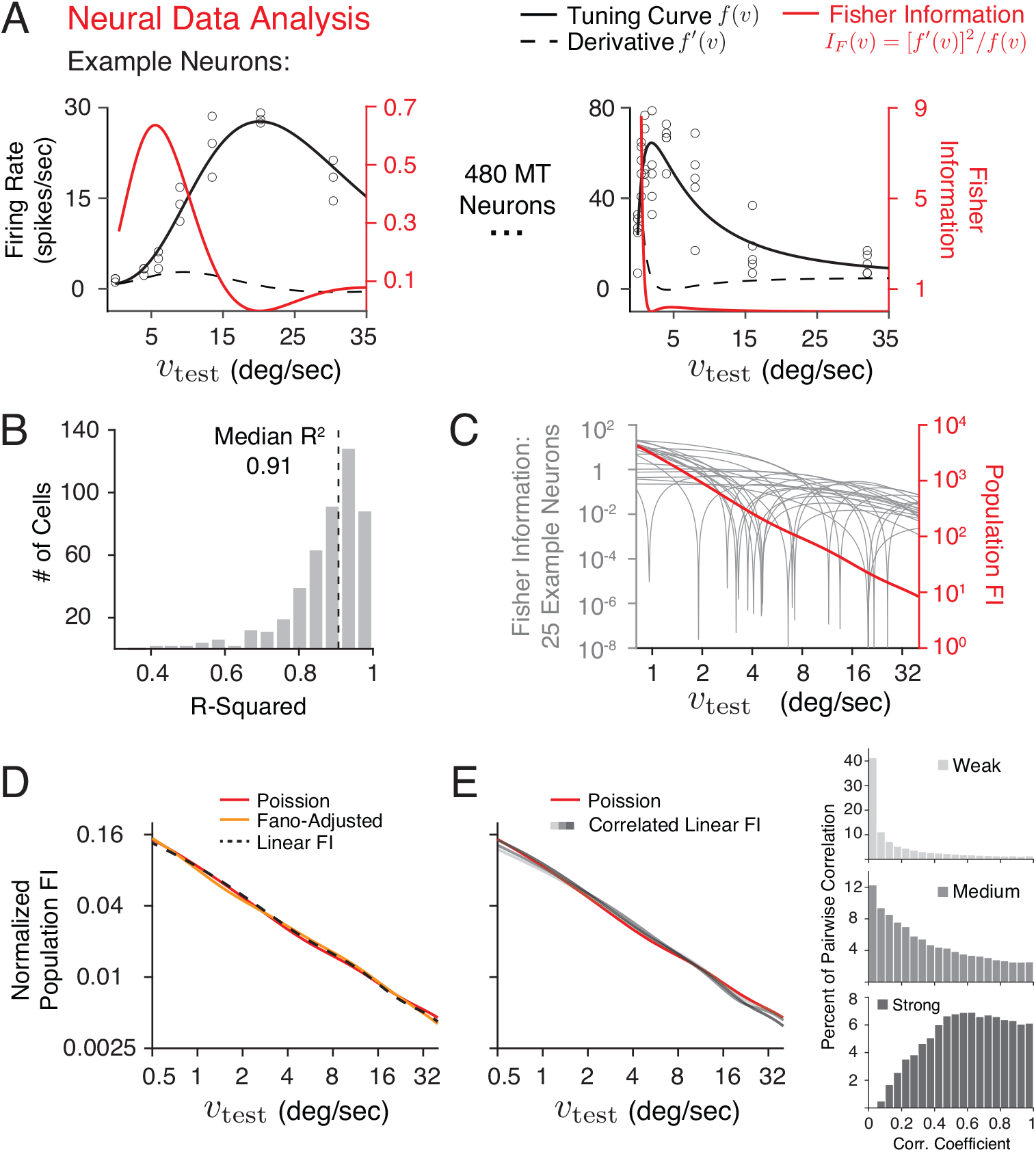
Extracting the “neural prior”. According to the efficient coding constraint (Eq. (1)) the population Fisher information (FI) should directly reflect the prior distribution of visual speed. **A)** Mean firing rates as a function of stimulus speed *v*_test_ (dots) shown for two example MT neurons from the data set (Nover et al., 2005), together with their fit log-normal tuning curves (black, left y-axis) and corresponding FI (red, right y-axis) assuming a Poisson noise model. **B)** Histogram of the goodness of the log-normal tuning curve fit across all neurons measured by *R*^2^. **C)** Individual FI of 25 example neurons (gray, left y-axis), and the population FI (red, right y-axis), calculated as the sum of the FI over all 480 neurons in the data set. **D)** The normalized square-root of population FI assuming independent Poisson noise (red), adjusted for the variance by estimating the Fano factor explicitly (orange), and the linear Fisher information (dashed black). **E)** The normalized square-root of population FI assuming independent Poisson noise (red), and the linear population FI based on a limited-range correlation model (Abbott and Dayan, 1999) for three levels of correlation strength as illustrated by the histogram of pairwise correlation coefficients on the right. See *Methods* for details.

Given the inherent limitations of single-unit recorded data, assumptions about the noise and its correlation structure in the population are necessary in order to compute the population FI (Abbott and Dayan, 1999; Averbeck et al., 2006; Moreno-Bote et al., 2014; Kohn et al., 2016). We first considered the noise to be independent among the neurons in the recorded population. In this simplified scenario, the population FI is the sum of the FI of individual neurons (Fig. 7C). The shape of the resulting population FI is very close to a power-law function; that is, when plotted on a log-log scale it closely resembles a straight line. Two slightly different methods of calculating the FI of individual neurons, either by estimating the Fano factor explicitly or by computing the linear FI (Kanitscheider et al., 2015; Kohn et al., 2016), produced nearly identical estimates of the shape of the population FI (Fig. 7D; *Methods*).

Further, assuming a correlation pattern that is speed-independent simply reduces the magnitude of the population FI but does not change its overall shape compared to the independent noise assumption. However, tuning-dependent noise correlations between pairs of neurons have been reported for area MT (Huang and Lisberger, 2009). Thus, to assess the potential impact of such correlations, we computed the linear population FI with a limited-range correlation model based on the relative speed preferences of individual neurons (Abbott and Dayan, 1999) (*Methods*). We found that although the magnitude of FI decreases with increasing correlation strength, the shape of the population FI is largely invariant within a large range of simulated correlation strengths (Fig. 7E). The reason why these correlations have little effect on the shape of the population FI is that the tuning characteristics of MT neurons are relatively “homogeneous” (i.e., the parameters of the tuning curve, such as the tuning width, are mostly independent of speed preference), and close to uniformly tile the logarithmic speed space (Nover et al., 2005). Thus, we argue that given the available evidence, estimating the *shape* of the population FI assuming independent noise is a reliable approximation.

The efficient coding constraint makes the additional prediction that the overall magnitude of the population FI corresponds to the total represented sensory information, and thus should be directly related to the contrast-dependent noise parameter of our observer model (Wei and Stocker, 2015, 2016; Noel et al., 2021). Although the fit noise parameter values are consistent with the typical contrast response function of cortical neurons (Fig. 3), a rigorous test of the prediction requires characterization of MT speed encoding at different levels of stimulus contrasts, which is something the current data do not provide. Preliminary neural data (Stocker et al., 2009) suggest, however, that stimulus contrast indeed simply scales the population FI without changing its shape. This is intriguing given the well-documented diversity and heterogeneity by which stimulus contrast affects the shape and position of speed tuning curves in area MT (Pack et al., 2005; Krekelberg et al., 2006; Stocker et al., 2009).

### Comparing the behavior and neural prior

Finally, we compared the extracted behavioral and neural priors. If our observer model is correct then the prior expectation with which a subject perceives the speed of a moving stimulus should be quantitatively identical to, and thus predictive of, the stimulus distribution for which neural encoding is optimized. Figure 8A shows the extracted behavioral prior of every subject and the neural prior. The prior distributions are indeed very similar and are consistent with a power-law function with an exponent of approximately −1.

**Figure 8:**
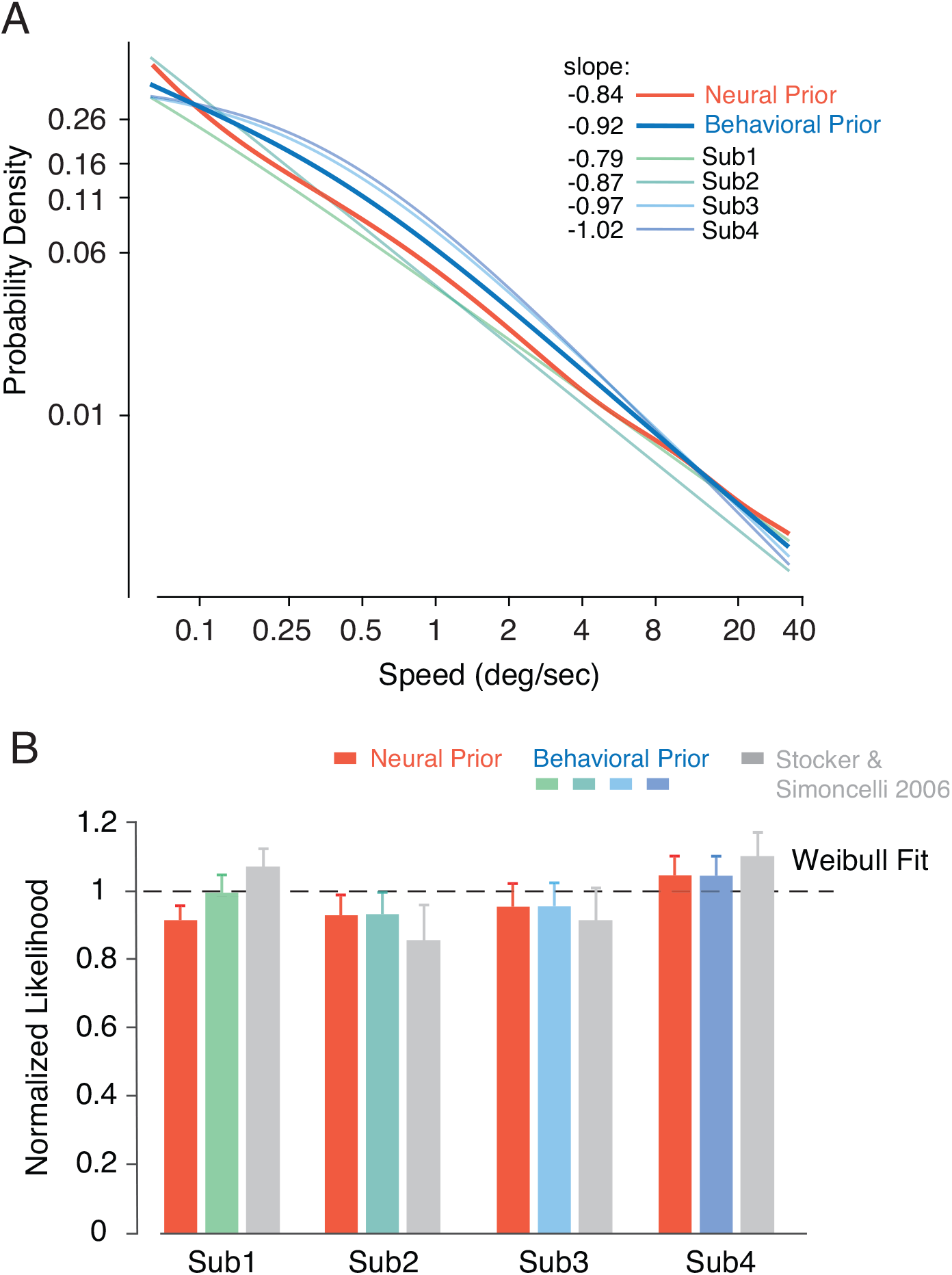
Comparing neural and behavioral prior. **A)** Reverse-engineered behavioral priors of every subject and their average (dark blue) superimposed by the neural prior (red). Slope values are computed from a linear fit of the curves in log-log coordinates. **B)** The cross-validated log-likelihood of the model using the subject’s best-fitting behavioral prior (blue) or the fixed neural prior (red), and the log-likelihood of the original model (Stocker and Simoncelli, 2006) (gray). The log-likelihood value is normalized to the range defined by a “coin-flip” model (lower bound) and the Weibull fit to each psychometric curve (upper bound). Error bars represent ± SD across 100 validation runs according to a five-fold cross-validation procedure. See *Methods* for details.

In order to quantitatively assess the effective similarity between the behavior and neural prior, we constructed a “neural observer” model for which the prior was fixed to be the neural prior extracted from the MT data; only the contrast-dependent noise parameters *h*(*c*) were free parameters. We used a cross-validation procedure to compare this neural observer with the unconstrained observer model and the original model (Stocker and Simoncelli, 2006). As illustrated in Fig. 8B, the validation performances are highly similar across all three models and closely match the performance of individual Weibull fits. This comparison demonstrates several aspects. First, it confirms that the behavioral and neural prior are behaviorally indistinguishable and thus effectively equivalent; if the neural data did not exist, we would have been able to accurately predict the encoding accuracy of MT neurons at the population level. Second, it highlights the excellent quality of the Bayesian observer model as its account of human behavior is close to that of the best possible parametric description of the data (i.e. individual Weibull fits). And finally, it shows that the complexity of our new observer model is appropriate and does not lead to over-fitting.

### Weber-Fechner law

The power-law shape of the behavioral and neural prior distribution also sheds a new normative light on the interpretation of Weber’s law. Famously, Fechner proposed that Weber’s law emerges from a logarithmic neural encoding of a stimulus variable (Fechner, 1860). Neural encoding of visual speed in area MT is indeed considered logarithmic (Nover et al., 2005; Burge and Geisler, 2015): When analyzed in the logarithmic speed domain, the tuning curves of MT neurons are approximately bell-shaped, scale-invariant, and tile the stimulus space with near-uniform density (Nover et al., 2005; Pack et al., 2005).

With our new model, we have already demonstrated that a modified power-law prior (Eq. (2)) with an exponent of approximately −1 can well account for Weber’s law behavior and the deviation from it at slow speeds (Fig. 6). We can now show that this power-law prior also predicts logarithmic neural encoding. Specifically, one way to implement the efficient coding constraint (Eq. (1)) is to assume a homogeneous neural encoding (i.e., identical tuning curves that uniformly tile the sensory space) of the variable of interest *transformed* by its cumulative distribution function (CDF) (Ganguli and Simoncelli, 2010; Wei and Stocker, 2012; Wang et al., 2016), which is sometimes also referred to as histogram equalization (Acharya and Ray, 2005). With an exponent *c*_0_ = −1, the CDF of the speed prior is exactly the logarithmic function that well-described MT tuning characteristics (Nover et al., 2005) (*Methods*). Thus, the Bayesian observer model constrained by efficient coding provides a normative explanation for both Weber’s law and the logarithmic encoding of visual speed in area MT.

## Discussion

We presented a Bayesian observer model constrained by efficient coding for human visual speed perception. We fit this model to existing human 2AFC speed discrimination data recorded over a wide range of stimulus contrasts and speeds, which allowed us to reverse-engineer the “behavior prior” that best accounts for the psychophysical behavior of individual subjects. In addition, we analyzed the population encoding accuracy of visual speed based on an existing set of single-cell recordings in area MT, thereby extracting the “neural prior” according to the efficient coding constraint of our observer model. We found that the behavioral prior estimated from the psychophysical data accurately predicts the neural prior reflected in the encoding characteristics of the MT neural population.

Our results provide a successful, quantitative validation of the Bayesian observer model constrained by efficient coding in the domain of visual speed perception. We demonstrate that this model can accurately account for the behavioral characteristics of bias and threshold in visual speed perception if subjects’ prior belief about the statistical distribution of visual speed resembles a power-law function with an exponent of approximately −1. Cross-validation revealed no significant difference between the best possible parametric description of the behavioral data (i.e., individual Weibull fits) and our model fits. Compared to the original, more flexible Bayesian model formulation (Stocker and Simoncelli, 2006), the added efficient coding constraint results in estimates of behavioral priors that are not only much more consistent across subjects but also remarkably predictive of the population encoding characteristics of neurons in the motion-sensitive area MT in the primate brain. Our work substantially strengthens the evidence for the slow-speed prior interpretation of motion illusions (Weiss et al., 2002; Stocker and Simoncelli, 2004, 2006; Welchman et al., 2008; Sotiropoulos et al., 2014; Jogan and Stocker, 2015; Senna et al., 2015) (but see (Rideaux and Welchman, 2020)) by the demonstrated quantitative support from electrophysiological neural data.

We also offer an explanation for why certain perceptual variables have a logarithmic neural representation and thus follow Weber’s law (Fechner, 1860). According to our model, logarithmic encoding and Weber’s law both follow from the efficient representation of a perceptual variable with a power-law prior distribution. We thus predict that perceptual variables that conform to Weber’s law have power-law distributions with an exponent of approximately −1 *and* are logarithmically encoded in the brain (although alternative encoding solutions that satisfy the efficient coding constraint are possible, see (e.g. Wei and Stocker, 2015)). Indeed, perceptual variables that are known to approximately follow Weber’s law such as weight (Fechner et al., 1966), light intensity (Treisman, 1964), and numerosity (Nieder and Miller, 2003; Cheyette and Piantadosi, 2020; Prat-Carrabin and Woodford, 2021), exhibit heavy tails in their statistical distributions under natural environmental conditions (Dehaene and Mehler, 1992; Dror et al., 2004; Peters et al., 2015; Piantadosi and Cantlon, 2017), a defining feature of a power-law function. Conversely, any deviation from Weber’s law and the logarithmic encoding should be reflected in deviations of the statistical stimulus distributions from a power-law function. Future studies of natural stimulus statistics, modeling of psychophysical data, and neural recordings will be needed to further and more quantitatively validate the generality of this prediction.

Other recent work has used efficient coding assumptions to link perceptual discriminability to the statistical prior distribution of perceptual variables (Gu et al., 2010; Ganguli and Simoncelli, 2016; Sims, 2018). Our approach is a substantial step forward in that it embeds this link within a full behavioral observer model. Thus rather than relying on single summary metric of behavior (i.e. discrimination threshold), the predictions of our model are constrained by the full richness of the psychophysical data, i.e., every single datum in the set. This not only provides a much more stringent test of the observer model but also permits more robust and precise predictions of neural coding accuracy and priors.

The presented comparison between behavioral and neural prior is limited to the extent that there were substantial experimental differences between the behavioral and neural data. For example, we compared human with non-human primate data and estimated the neural tuning characterization based on single-cell responses to random-dot motion rather than the broadband, drifting grating stimuli used in the psychophysical experiment. Yet, the surprisingly accurate match of the extracted neural and behavioral priors suggests that they may reflect the “true” stimulus prior, in which case these differences in stimuli and model systems should indeed matter little because the stimulus prior is largely a property of the environment and not the observer nor the particular stimulus pattern. Recent studies have demonstrated that it is possible to quantitatively characterize the accuracy with which a perceptual variable is represented in the human brain using voxel-level encoding models of functional magnetic resonance imaging signals (Van Bergen et al., 2015). Future work may exploit this technique to validate and potentially refine our estimates of the “neural prior” in human subjects. Such work would also permit a more thorough investigation of individual differences at both behavioral and neural levels through matched task and stimulus designs.

The specific shape of the extracted neural and behavioral prior depends on the assumed efficient coding objective. The chosen efficient coding constraint (Eq. (1)) results from the objective to maximize the mutual information between neural representation and stimulus (Wei and Stocker, 2016). It is possible, although unlikely given the exceptional good quantitative match, that with a different combination of efficient coding constraint and loss function (see Wang et al., 2012; Morais and Pillow, 2018; Rast and Drugowitsch, 2020), a power-law prior with a different exponent could also be consistent with both the behavioral and neural data. This is difficult to validate conclusively without access to an accurate characterization of the stimulus prior of visual speed, as the search space over all possible combinations is extensive. Previous work has shown, however, that the encoding characteristics in early visual cortex for visual stimulus variables for which good estimates of the stimulus prior exist (e.g., luminance contrast and local orientation) are closely accounted for by the mutual information maximization objective (Wang et al., 2012).

An important assumption of our observer model is that the neural and behavioral priors not only match but are also consistent with the statistical distribution of visual speeds in the natural environment (“stimulus prior”). As such, our results predict that the stimulus prior approximates a power-law distribution that lies within the range given by the neural and behavioral priors shown in Fig. 8A. However, an empirical validation of this prediction by directly measuring the stimulus prior distribution is rather challenging. Object motion, but also the ego-motion of the observer in terms of its body, head, and eye movements all contribute to the visual motion signal. Thus, the precise characterization of the visual speed distribution would require accurate measurements and calibrations of these different types of motions, as well as of the algorithm used to extract the motion information from the visual signal. Previous studies have approximated these relative movements to various degrees and used different algorithms to extract local visual speed from spatio-temporal images, resulting in different characterization of the prior distribution (Roth and Black, 2007; Baker et al., 2011; Sinha et al., 2021; Dong and Atick, 1995). However, common to all these measured stimulus priors is that they have higher probabilities at slow speeds and form long-tailed distributions. Future work using more comprehensive data (DuTell et al., 2020) may provide a better characterization of visual speed priors under ecologically valid, natural conditions.

We expect our model and analytic approach to be applicable to any other perceptual variable and task that exhibit characteristic patterns of perceptual biases and discrimination thresholds. However, of particular interest and posing a strong test of our model are changes in perceptual bias and threshold that are induced by spatio-temporal context such as adaptation aftereffects or the tilt-illusion (Schwartz et al., 2007, 2009; Clifford et al., 2007). It is traditionally assumed that these biases are caused by a mismatch in expectation between encoding and decoding (i.e. the “coding catastrophe” (Schwartz et al., 2007)), which is in sharp contrast to one of the main features of our model. Preliminary results are promising (Wei and Stocker, 2017; Wei et al., 2015). However, more quantitative analyses are necessary to test how well the framework can account for the data and what neural and behavioral priors it will predict.

In summary, within the context of visual speed perception, we have demonstrated that the Bayesian observer model constrained by efficient coding has the potential to provide a unifying framework that can quantitatively link natural scene statistics with psychophysical behavior and neural representation. Our results represent a rare example in cognitive science where behavioral and neural data *quantitatively* match within the predictions of a normative theory.

## Acknowledgments

We thank Greg DeAngelis for sharing his electrophysiological MT data, Benjamin Chin for his work on an earlier implementation of the model, Emily Cooper, Mike Landy and Rafael Polania for their helpful comments on the manuscript, and the members of the Computational Perception and Cognition Laboratory for many fruitful discussions of the work.

## Extended Data Figures

**Extended Data Figure 2 - 1:**
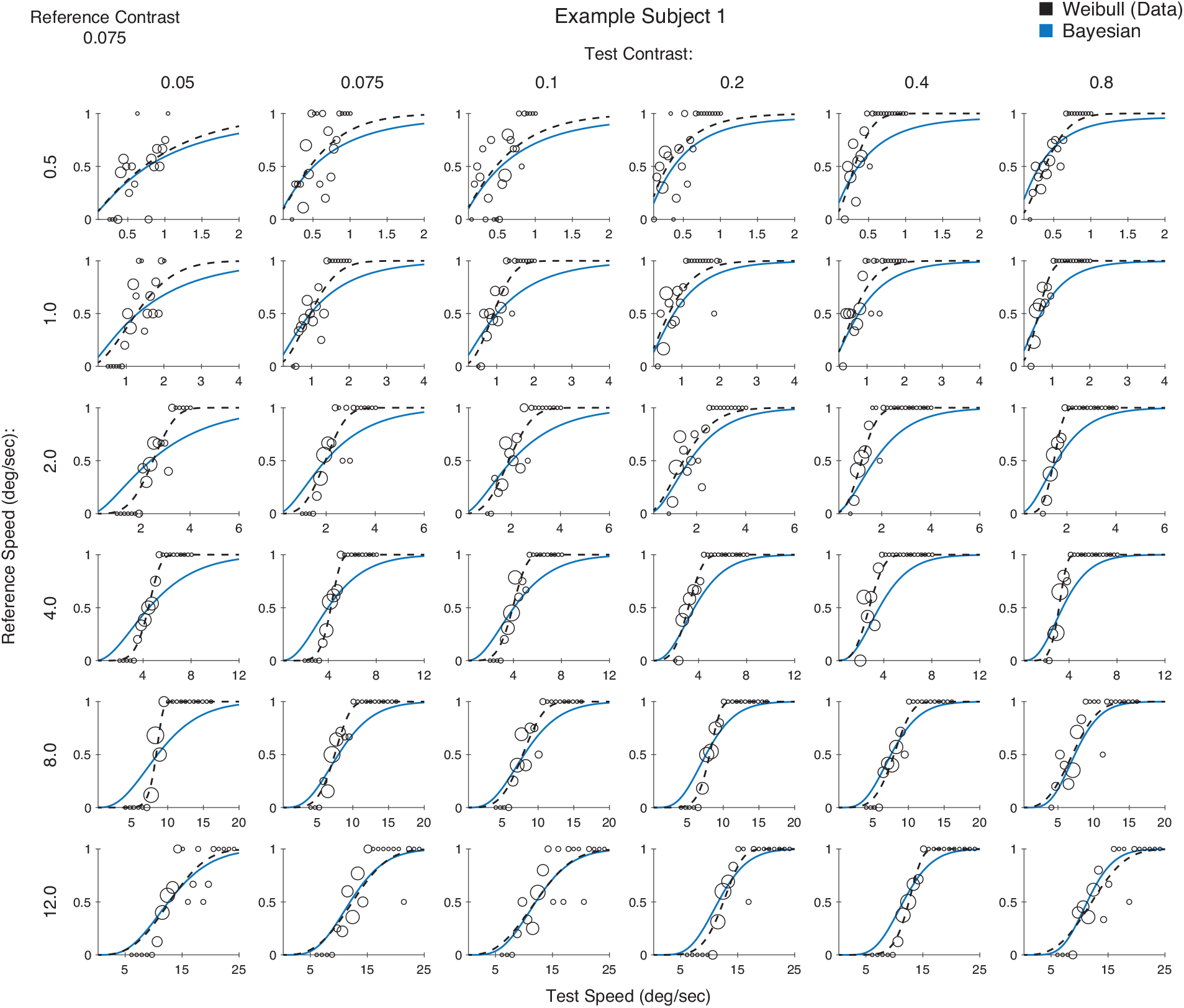
Psychometric curves across all conditions for exemplary Subject 1 (low-contrast reference). Contrast of the reference stimulus was 0.075. Each row corresponds to a different reference speed as indicated on the left, and each column corresponds to a different test contrast as indicated on top. The dashed black curve represents a Weibull fit to the data of each individual condition, and the blue curve is the Bayesian model fit to the entire data set. The size of the circle is proportional to the number of trials at that value.

**Extended Data Figure 2 - 2:**
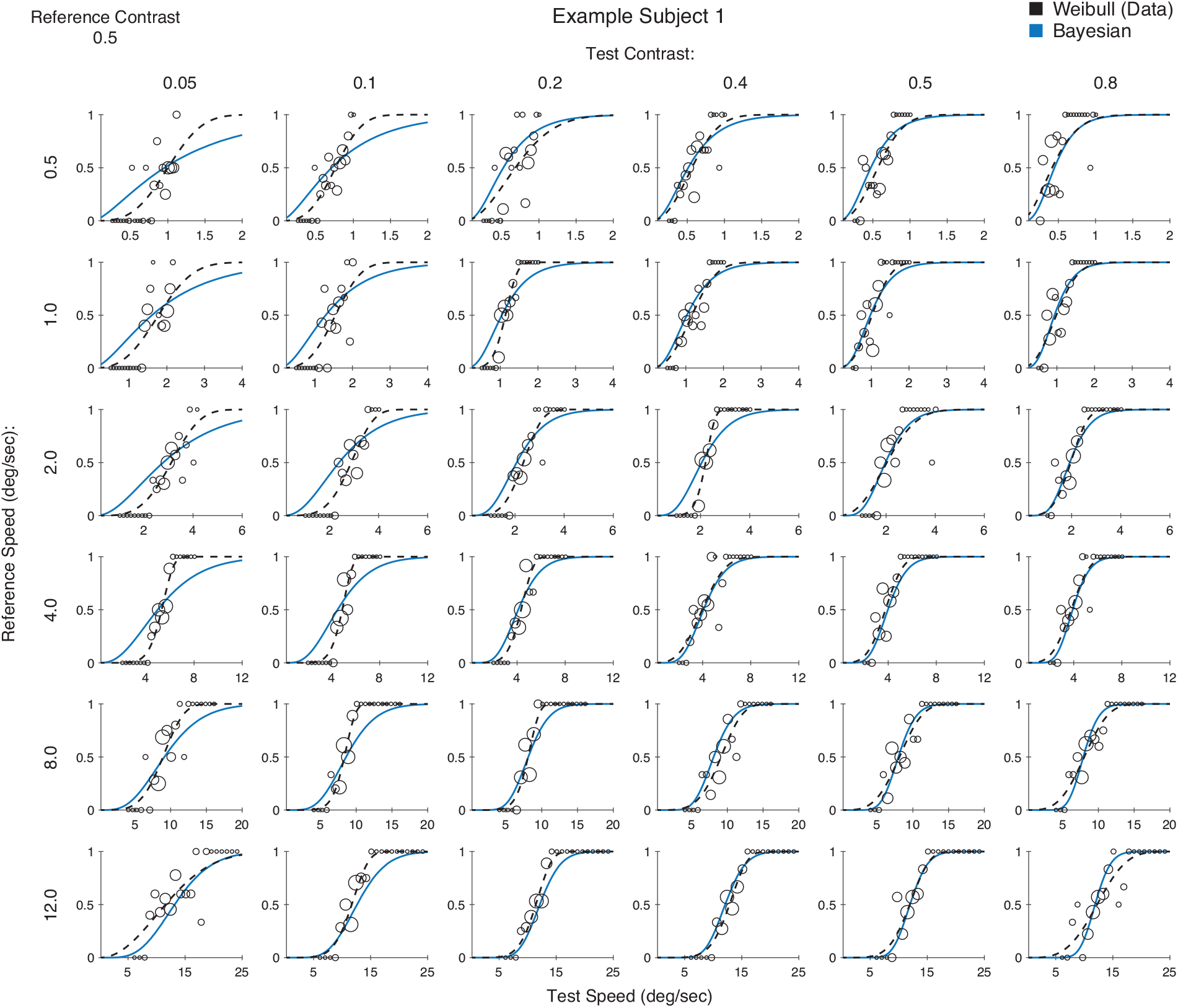
Psychometric curves across all conditions for exemplary subject 1 (high-contrast reference). Contrast of the reference stimulus was 0.5. Each row corresponds to a different reference speed as indicated on the left, and each column corresponds to a different test contrast as indicated on top. The dashed black curve represents a Weibull fit to the data of each individual condition, and the blue curve is the Bayesian model fit to the entire data set. The size of the circle is proportional to the number of trials at that value.

